# Evaluation of the effects of ripening-related genes on fruit metabolites and the associated regulatory mechanisms in tomato

**DOI:** 10.1101/2024.01.08.574695

**Authors:** Huimin Jia, Yaping Xu, Yuanwen Deng, Yinhuan Xie, Zhongshan Gao, Zhaobo Lang, Qingfeng Niu

**Affiliations:** College of Agronomy, Jiangxi Agricultural University, Nanchang 330045, China; Advanced Academy, Anhui Agricultural University, Research Centre for Biological Breeding Technology, Hefei, Anhui, 230036, China; National Engineering Laboratory of Crop Stress Resistance Breeding, School of Life Sciences, Anhui Agricultural University, Hefei, 230036, China; Department of Horticulture, College of Agriculture and Biotechnology, Zhejiang University, 310058, Hangzhou, Zhejiang, China; Institute of Advanced Biotechnology, School of Life Sciences, Southern University of Science and Technology, Shenzhen 518055, China

**Author notes:** Co-corresponding authors: Zhaobo Lang and Qingfeng Niu. Co-first authors (Huimin Jia, Yaping Xu).

**Keywords:** Ripening-related TFs, Metabolites, Transcriptome, Sugar accumulation, *HY5*, *SWEET12c*

## Abstract

Fruit ripening, which is a complex process involving dynamic changes to metabolites, is controlled by multiple factors, including transcription factors (TFs). Several TFs are reportedly essential regulators of tomato fruit ripening. To evaluate the effects of specific TFs on metabolite accumulation during fruit ripening, CRISPR/Cas9-mediated mutagenesis was combined with metabolome and transcriptome analyses to explore regulatory mechanisms. Specifically, we generated various genetically engineered tomato lines that differed regarding metabolite contents and fruit colors. The metabolite and transcript profiles indicated that the selected TFs have distinct functions that control fruit metabolite contents, especially carotenoids and sugars. Our findings may provide new insights into the regulatory mechanisms governing tomato fruit ripening. Moreover, a mutation to *ELONGATED HYPOCOTYL5* (*HY5*) increased the tomato fruit fructose and glucose contents by approximately 20% (relative to the wild-type levels). Our *in vitro* assay showed that HY5 can bind directly to the G-box *cis*-element in the *SWEET12c* promoter to activate expression, thereby modulating sugar transport. Our findings have clarified the mechanism regulating fruit metabolic networks, while also providing the theoretical basis for breeding horticultural crops that produce fruit with diverse flavors and colors.g

## Introduction

Tomato is one of the most important vegetable crops worldwide (León-García et al., 2017). It is also a model plant for investigating climacteric fruit ripening because of its short life cycle and relatively small genome (approximately 780 Mb) that is easy to modify (Alexander and Grierson, 2002; Sato et al., 2012). The aroma-related volatiles, fruit flavors, and characteristic pigments typically produced during the fruit ripening period are important for breeding and genetic selection (Carrari et al., 2006). Plant metabolites, such as carotenoids, flavonoids, and sugars, undergo dramatic changes during the softening of fruit in the ripening transition stage (Li et al., 2020), making them important indicators of fruit maturity and quality. The metabolic changes that occur as tomato fruit ripen are controlled by phytohormones (mainly ethylene) and numerous ripening-related genes (RRGs) (Zhong et al., 2013; Liu et al., 2015). Over the past 40 years, there have been considerable advances in tomato genetics research. For example, many spontaneous fruit ripening-related mutants have been identified under natural conditions, including the following: *ripening-inhibitor* (*rin*), *non-ripening* (*nor*), *Colorless non-ripening* (*Cnr*), *Never-ripe* (*Nr*), and *Green-ripe* (*Gr*). The fruit of the *rin*, *nor*, and *Cnr* mutants fail to ripen (Klee and Giovannoni, 2011; Li et al., 2019). The *rin* locus encodes a fusion protein comprising the truncated MADS-box transcription factors (TFs) RIPENING INHIBITOR (RIN) and MC. The *RIN* gene encodes SEPALATA (SEP), which combines with the MADS-box protein AGAMOUS-LIKE1 (TAGL1) or the two FRUITFULL homologs TDR4 (also called FUL1) and MBP7 (also called FUL2) to form complexes that regulate ripening (Itkin et al., 2009; Martel et al., 2011). Additionally, RIN is a master positive regulator that is critical for ripening (Li et al., 2019) because it targets RRGs affecting metabolic pathways, including genes encoding proteins associated with ethylene and carotenoids [phytoene synthase 1 (PSY1) and ζ-carotene desaturase (ZDS)] as well as texture [polygalacturonase 2a (PG2a), zeta-carotene isomerase 4 (TBG4), expansin 1 (EXP1), and endo-β-1,4-glucanase (CEL2)], flavor, aroma, and sucrose metabolism (Martel et al., 2011; Li et al., 2020).

Tomato is a useful system for studying the relationship between ethylene and ripening-related factors. Several regulators that substantially affect the initial ripening period, including RIN, TDR4, MBP7, NOR, TAGL1, APETALA2a (AP2a), and HY5, function in ethylene-dependent or ethylene-independent pathways that modulate fruit maturation (Itkin et al., 2009; Chung et al., 2010; Bemer et al., 2012; Gao et al., 2020). On the basis of the lack of ripening resulting from spontaneous mutations, RIN, NOR, and CNR were long considered to be major regulators required for inducing ripening. However, recent studies involving knockout mutants generated by the CRISPR/Cas9 system revealed that the regulation of fruit ripening is not solely dependent on these ripening-related TFs (Ito et al., 2017; Gao et al., 2019; Gao et al., 2020). There is evidence that RIN (Ito et al., 2017; Li et al., 2020), NOR (Gao et al., 2020), TDR4, CNR, and AP2a (Wang et al., 2019) function downstream of the ethylene signaling pathway and form a transcriptional feedback circuit with ETHYLENE INSENSITIVE3-Like (EIL) proteins to precisely control autocatalytic ethylene production during ripening (Lü et al., 2018; Li et al., 2019; Huang et al., 2022; Deng et al., 2023). Therefore, the functions of ripening-related TFs during ripening must be re-evaluated and the relationship between these regulators and the ethylene-dependent or ethylene-independent pathway should be clarified. In several studies, genomes were edited to generate mutants and then metabolomics and transcriptomics techniques were applied to analyze gene functions and effects on metabolites. For example, HY5 reportedly promotes fruit ripening and chlorophyll accumulation. In the *Slhy5* mutant, the fruit exhibit delayed ripening, have a relatively low chlorophyll content during the mature green fruit stage, and appear light red in the final ripening stage. Metabolome and transcriptome analyses of the fruit indicated that HY5 positively regulates the accumulation of flavonoids and caffeoylquinic acid (Zhang et al., 2022). The *NOR-like1* gene, which encodes a TF belonging to the NAC family, is a newly identified positive regulator of tomato fruit ripening. Using CRISPR/Cas9 technology to target this gene resulted in the production of mutant fruit with decreased lycopene accumulation and ethylene production, but increased alkaloid contents compared with the corresponding wild-type (WT) levels. Moreover, NOR-like1 also affects phenolic acids and flavonoids (Yang et al., 2022). Although the effects of some TFs on fruit metabolism have been determined, many TFs remain uncharacterized and there is a lack of systematic research on how fruit ripening-related TFs modulate fruit metabolism.

In this study, we investigated the major regulatory effects of TFs on tomato fruit metabolite compositions. Specifically, we analyzed the effects of knocking out *AP2a*, *HY5*, *RIN*, *TAGL1*, *TDR4*, *MBP7*, *RIN/TDR4*, *NOR*, and *NR* using CRISPR/Cas9 technology on ‘Ailsa Craig’ (AC) tomato fruit, while also performing metabolomics and transcriptome sequencing (RNA-seq) analyses. According to our results, mutations to the selected TF genes adversely altered the normal fruit ripening process by modifying pigment accumulation, ethylene release, or other fruit ripening-related factors. The mutations to certain genes created mutants with a visually similar phenotype, but diverse carotenoid contents (e.g., *MBP7* and *TAGL1*; *RIN*, *NOR*, and *AP2a*). The TF mutations differentially impaired the accumulation of phenolic compounds, linoleic acids, and steroidal glycoalkaloids (SGAs) because of changes to the expression of target genes involved in pathways mediating biosynthesis and metabolism. The accumulation of SGAs depended on ethylene and influenced ripening. Earlier research indicated RIN is crucial for ripening because it regulates ethylene production (Iijima et al., 2009; Kazachkova et al., 2021). The SGA contents were higher in the *rin-cr1*, *tdr-cr1*, and *mbp7-cr1* mutant fruit than in the AC fruit, implying RIN and its cofactors promote the conversion of SGAs to less toxic metabolites. A mutation to *HY5* decreased flavonoid and linoleic acid contents. Furthermore, the SGAs accumulated similarly between *hy5-cr1* and AC. Surprisingly, the fructose and glucose contents increased by approximately 20% in the *hy5* fruit. Further analyses suggested that HY5 represses the accumulation of sugar in tomato fruit by activating *SWEET12c* expression. This study provides new insights into how TFs affect the tomato metabolome. The data provided herein may be useful for breeding tomato varieties with ideal metabolite compositions.

## Results and Discussion

### Ripening*-*related TFs alter the tomato fruit carotenoid content via pathways dependent or independent of ethylene

Recent studies have used CRISPR/Cas9 gene-editing technology to analyze gene functions in plants (Ito et al., 2017; Wang et al., 2019; Gao et al., 2020). In this study, we mutated specific genes (*HY5*, *AP2a*, *TAGL1*, *MBP7*, *RIN*, *RIN/TDR4*, *TDR4*, *NOR*, and *NR*) using this gene-editing system to clarify how the encoded proteins regulate fruit ripening and metabolite contents (Supplementary Figure S1). Because the mutant lines for each gene were very similar, we selected only one mutant line per gene (i.e., *ap2a-cr1*, *hy5-cr1*, *tagl1-cr1*, *mbp7-cr1*, *rin-cr1*, *tdr4-cr1*, *rin/tdr4-cr1*, *nor-cr1*, and *nr-cr1*) for further analyses. Diverse fruit colors were observed among the mutants during the ripening stage (Fig. 1A). Carotenoids, which are the main pigments responsible for tomato fruit colors (generally yellow, orange, and red), are naturally occurring pigments with beneficial effects on human health (Liu et al., 2015). To gain some insights into the underlying causes of the fruit color differences between the nine mutants and AC at 46 days post-anthesis (DPA), the fruit carotenoid components and contents were analyzed using a UPLC-MS/MS system. A total of 42 carotenoids were identified in tomato fruit, including six carotenes and 36 xanthophylls (Supplementary Table S1), among which (E/Z)-phytoene, lycopene, lutein, and β- carotene were the main components in ripe tomato fruit (90.56% of the total carotenoid content) (Fig. 1B and Supplementary Figure S2). The carotenoid contents among the mutants ranged from 95.36 μg/g fresh weight (FW) (*rin/tdr4-cr1*) to 531.7 μg/g FW (*tagl1-cr1*). With the exception of *tagl1-cr1*, the mutants had a lower carotenoid content than the WT (446.90 μg/g FW), suggestive of the regulatory effects of the corresponding TFs on carotenoid biosynthesis. A previous study showed that the phenotype of *rin* was similar to that of *tagl1* and *nor* (Giovannoni et al., 2017). In the current study, the fruit of the *rin-cr1*, *tagl1-cr1*, and *nor-cr1* mutants were phenotypically similar, but there were significant differences in the carotenoid contents (Fig. 1B). The suppressed expression of *TAGL1* reportedly results in orange– yellow tomato fruit, with a decrease in the carotenoid content and an increase in the lutein content (Itkin et al., 2009). In the current study, the *tagl1-cr1* fruit were orange and had yellow stripes, likely because they had significantly higher lutein (2.24-fold) and β-carotene (1.46-fold) contents than the WT fruit (Fig. 1B). Although the carotenoid contents decreased by 10% in *ap2a-cr1* (402.3 μg/g FW) and by 7% in *mbp7-cr1* (415.9 μg/g FW), their fruit were completely red before turning orange– brown and red with stripes, respectively, when they were ripe (Wang et al., 2019). In addition, compared with the corresponding levels in the WT samples, the β-carotene and lutein contents were respectively 2.5-fold and 1.6-fold higher and the phytoene and lycopene contents were respectively 81.4% and 36.2% lower in *ap2a-cr1*. Accordingly, knocking out *AP2a* altered the fruit carotenoid components, which is in accordance with the findings of an earlier study in which decreased *AP2a* expression due to RNAi resulted in decreased carotenoid production (Chung et al., 2010). In *mbp7-cr1*, the β-carotene content increased by 1.33-fold (relative to the WT level), whereas the abundance of three other components decreased (10.8%–20%). Thus, the substantial decrease in the phytoene content likely explains the clear phenotypic difference between *ap2a-cr1* and the WT. The carotenoid content decreased by approximately 38.21%, 37.73%, 27.24%, and 25.08% in *tdr4-cr1*, *hy5-cr1*, *nor-cr1*, and *nr-cr1*, respectively, compared with the WT carotenoid content. Similar to the results of earlier research, the *nor-cr1* fruit examined in the present study were orange, with a decrease in the phytoene and lycopene contents (Wang et al., 2019). The orange-ripe phenotype of *tdr4-cr1* was similar to that of *TDR4* RNAi fruit (Bemer et al., 2012). The decrease in the contents of the four main carotenoids in *hy5-cr1* was consistent with the carotenoid content changes in CRISPR/Cas9 *hy5-cr1* mutant fruit (Zhang et al., 2022). The *rin/tdr4* double knockout mutants, which were analyzed for the first time in our study, had bright yellow fruit. The *rin-cr1* and *rin/tdr4-cr1* mutants had significantly lower carotenoid levels than the WT. Moreover, the phytoene and lycopene contents clearly decreased in the *rin-cr1* mutant and were undetectable in the double mutants (Li et al., 2020). To further examine how these TFs affect specific nutrients and fruit ripening, we analyzed ethylene production and performed metabolomics (UPLC-MS/MS) and RNA-seq analyses.

**Fig. 1.**
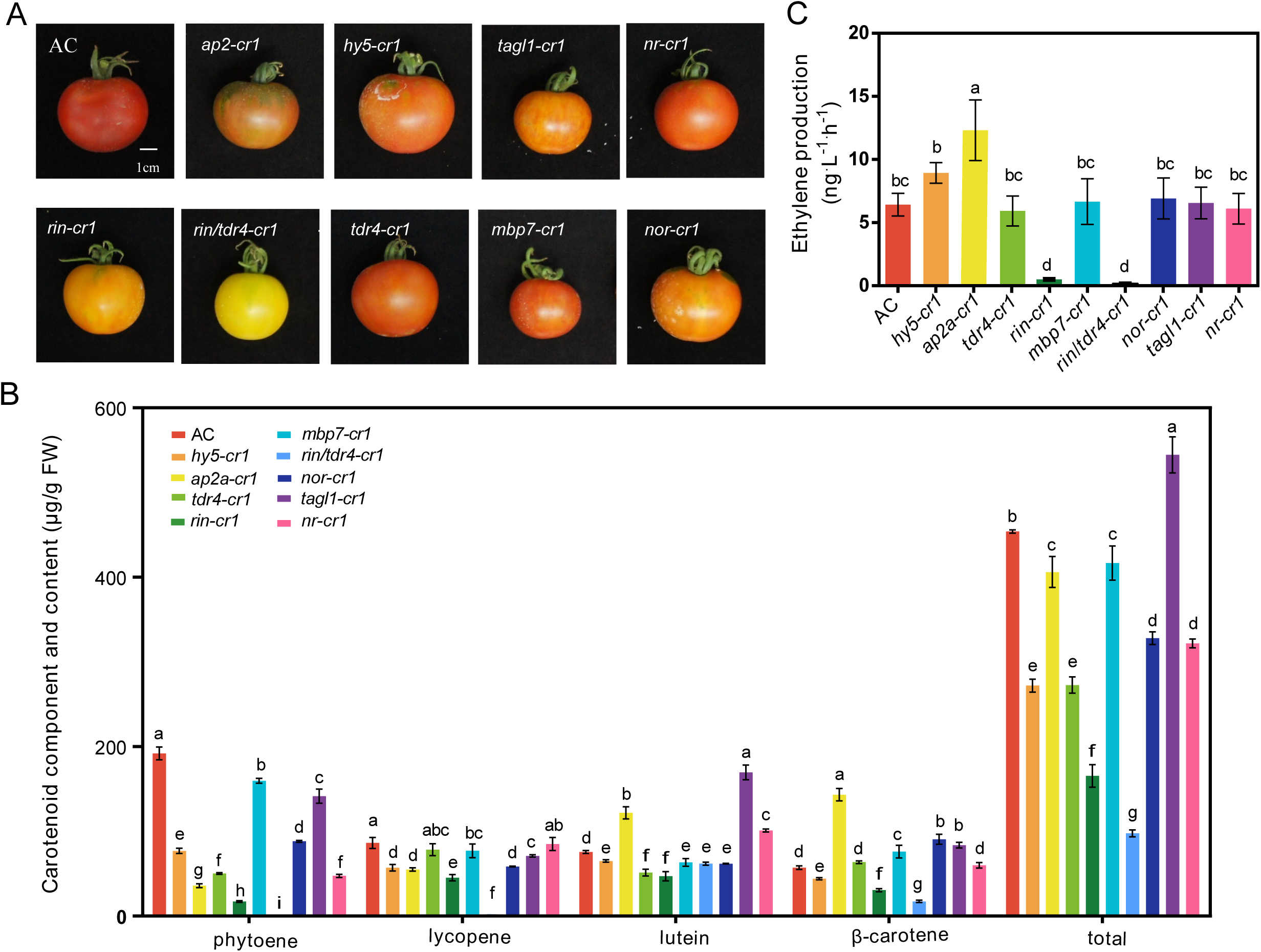
Phenotypes of mutant fruit and differences in the fruit ripening process between the mutants and ‘Ailsa Craig’ (AC). (A) Phenotypes of the mutant and AC fruit collected at 46 DPA. Scale bar, 1 cm. (B) Carotenoid contents of the mutants and AC. (C) Ethylene production in the mutants and AC at 46 DPA. Data are presented as the mean ± SD (n = 3) in panels B and C. Lowercase letters indicate significantly different values (P < 0.05).

Ethylene production in the fruit ripening stage was compared between the mutants and the WT (Fig. 1C). The 1.3-fold increase in ethylene production in *ap2a- cr1* (relative to the WT level) suggests that *AP2a* encodes a negative regulator of ethylene biosynthesis (Chung et al., 2010; Wang et al., 2019). The *rin-cr1* and *rin/tdr4-cr1* mutants contained a small amount of ethylene, implying *RIN* encodes a master regulator of ethylene production, whereas *TDR4* encodes a cofactor involved in ethylene biosynthesis (Li et al., 2020). The ethylene production in *hy5-cr1*, *tdr4-cr1*, *tagl1-cr1*, *nor-cr1*, and *nr-cr1* was similar to that in the WT, indicating these genes regulate fruit ripening independent or downstream of ethylene (Bemer et al., 2012). Considering the lack of a significant change to ethylene production in *nr-cr1*, *NR* may encode an ethylene receptor (ETR) that is functionally redundant with other ETRs (Tieman et al., 2000). Our data strongly indicate that RIN is a positive regulator of the carotenoid biosynthesis pathway along with the ripening process and ethylene burst. In addition, RIN and TDR4 cooperatively affect ripening, suggesting the regulatory effects of these TFs on fruit ripening involve ethylene as well as abscisic acid (ABA) or other hormones. Next, we screened for the carotenoid-regulating genes targeted by these TFs.

### The functional classification of the targeted genes revealed these TFs affect many biological processes associated with fruit ripening

To further functionally characterize the selected TFs in terms of their roles in the fruit ripening process, we analyzed the metabolomes and transcriptomes of the *ap2a-cr1*, *hy5-cr1*, *tagl1-cr1*, *mbp7-cr1*, *rin-cr1*, *rin*/*tdr4-cr1*, *nor-cr1*, *nr-cr1*, and AC tomato fruit at 46 DPA (red ripening stage). Both analyses included three biological replicates. The raw RNA-seq data are provided in Supplementary Table S2. According to the principal component analysis (PCA), the three replicates for each genotype were clustered together, reflecting the high reproducibility between biological replicates. Additionally, all samples were clearly separated into three distinct clusters (Fig. 2A). The *rin-cr1* and *rin/tdr4-cr1* mutants (in the upper left side of the plot) were separated from *mbp7-cr1* and *tdr4-cr1*, suggesting RIN might be the main component of the RIN-containing MADS-box complex that controls fruit ripening. The *hy5-cr1*, *ap2a-cr1*, *tagl1-cr1*, *nor-cr1*, and *nr-cr1* mutants as well as AC were clustered together, indicating the related downstream gene expression patterns differed slightly. The cluster dendrogram also included three distinct subgroups (Fig. 2A), which is consistent with the PCA data.

**Fig. 2.**
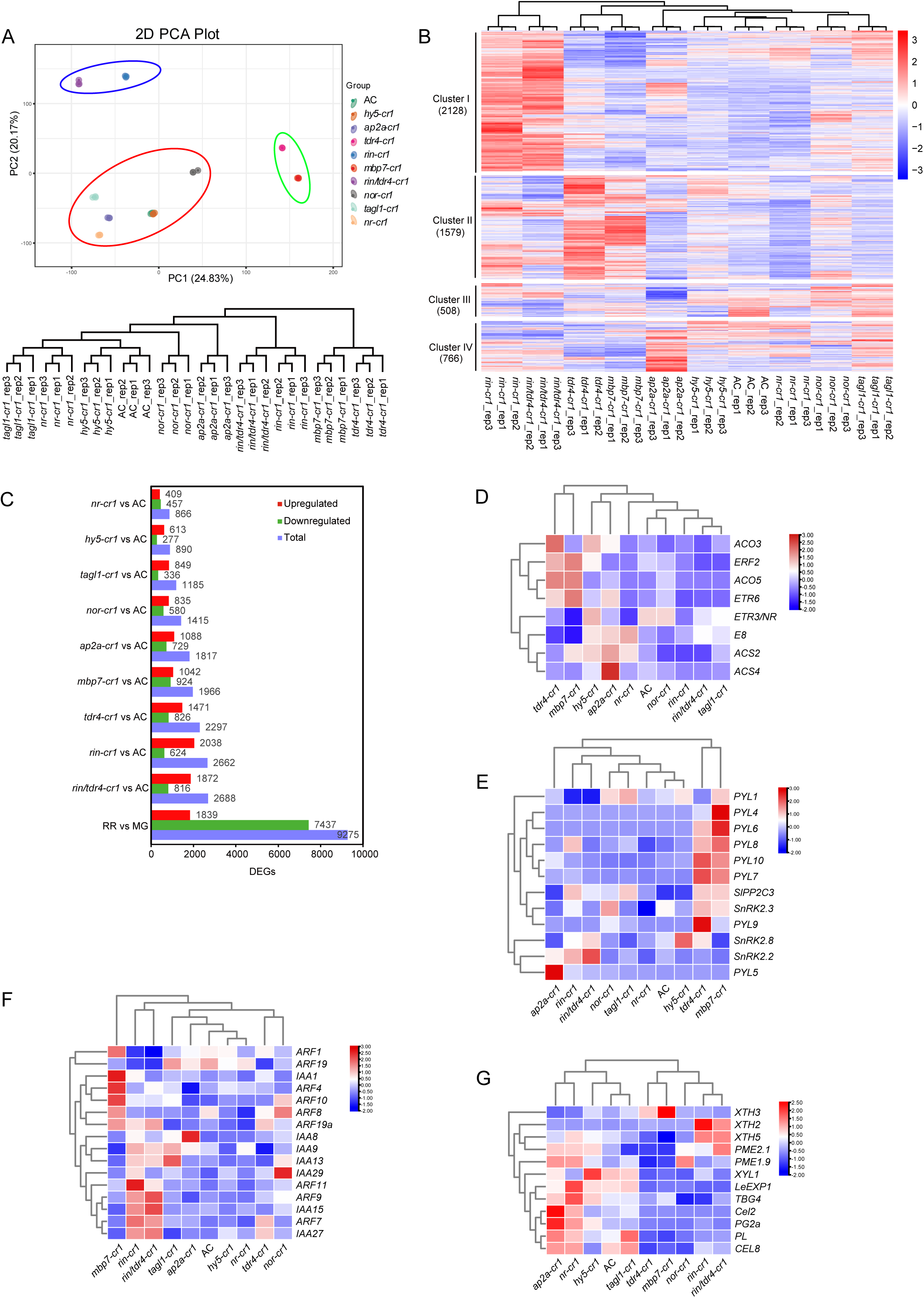
Genome-wide profiling of ripening-associated genes regulated by TFs. (A) Principal component analysis (PCA) plot and cluster dendrogram of the transcriptome data for the mutant and AC fruit. (B) Heatmap representation of the expression levels of the differentially expressed red ripening-related genes (RRGs) in the mutants and AC. (C) Distribution of the downregulated and upregulated genes among the TF-related differentially expressed genes (DEGs) and RRGs. Heatmaps were also constructed for the DEGs involved in ethylene biosynthesis and signal transduction (D), ABA signaling (E), IAA signaling (F), and cell wall metabolism (G).

To analyze the mechanism regulating fruit ripening in the nine mutants, the RRGs that were differentially expressed between the red ripening stage and the mature green stage were identified. A total of 9,275 RRGs were identified, including 1,839 upregulated and 7,437 downregulated genes (Supplementary Table S3). We next compared the transcript profiles of these RRGs between the mutant and AC fruit collected at 46 DPA (Fig. 2B, Supplementary Figure S4 and Tables S4–S12). There was a wide range in the transcript levels of RRGs in the fruit in the red ripening stage among the nine mutants and AC (Fig. 2B). A cluster analysis of the RRG expression patterns revealed four major clusters (Clusters I–IV) (Supplementary Table S13). The RRGs in these clusters were functionally annotated on the basis of a KEGG pathway analysis. The enriched KEGG pathways in these four clusters are summarized in Supplementary Figure S5. The cluster analysis detected distinct differences among these TFs. The significantly enriched terms were carbon metabolism (Cluster I), plant hormone signal transduction and phenylpropanoid biosynthesis (Cluster II), nucleotide and vitamin metabolism (Cluster III), and amino acid and terpenoid backbone biosynthesis (Cluster IV). The gene expression patterns were similar in the *nr-cr1*, *hy5-cr1*, *tagl1-cr1*, *nor-cr1*, and AC fruit (Fig. 2B and C). These results suggest that the mutations in these four mutants may be relevant to breeding because they delay fruit ripening without any noticeable large-scale changes in gene expression (relative to the expression in AC). Previous studies showed that genetically engineering *NOR* can result in tomato varieties that produce fruit with an extended shelf life (Kumar et al., 2018; Gao et al., 2020). Our results suggest that *HY5*, *NR*, and *TAGL1* should also be targeted by tomato breeders (i.e., silenced or genetically altered). The comparison with the WT control detected 866, 890, 1,185, 1,415, 1,817, 1,966, 2,297, 2,662, and 2,688 overlapping differentially expressed genes (DEGs) in the *nr-cr1*, *hy5-cr1*, *tagl1-cr1*, *nor-cr1*, *ap2a-cr1*, *mbp7-cr1*, *tdr4-cr1*, *rin-cr1*, and *rin/tdr4-cr1* mutants, respectively (Fig. 2C). The expression of more than half of the RRGs was influenced by eight specific TFs (Supplementary Figure S4B). These results imply the expression of numerous genes in ripening tomato fruit depends on TFs. Moreover, the *rin*/*tdr4-cr1 vs* AC and *rin-cr1 vs* AC comparisons detected more RRGs than the other comparisons. These findings further confirmed that RIN functions as a master activator that regulates the expression of hundreds of RRGs along with its cofactors TDR4 and MBP7 (Ito et al., 2017; Li et al., 2020).

Fruit ripening is controlled by a complex network comprising ripening-related regulators associated with ethylene, ABA, auxin, and secondary cell wall degradation. The expression patterns of ethylene biosynthesis and response genes (Fig. 2D), ABA signaling genes (Fig. 2E), auxin response genes (Fig. 2F), and cell wall-related genes (Fig. 2G) were examined. The ethylene production burst in fruit is predominantly attributed to the expression of ethylene biosynthetic genes encoding 1- aminocyclopropane-1-carboxylic acid synthase 2 (ACS2), ACS4, and 1- aminocyclopropane-1-carboxylic acid oxidase 1 (ACO1) (Barry and Giovannoni, 2007). In accordance with the observed ethylene production levels in the mutants, the *ACS2* and *ACS4* expression levels were downregulated in *rin-cr1* and *rin*/*tdr4-cr1*, but upregulated in *ap2a-cr1*. Accordingly, RIN may trigger the release of ethylene, whereas AP2a inhibits ethylene production, thereby controlling fruit ripening in a coordinated manner (Chung et al., 2010; Ito et al., 2017; Li et al., 2020). Notably, the expression levels of certain genes involved in ethylene signal transduction, including *ETHYLENE RESPONSE 6* (*ETR6*) and *ETHYLENE RESPONSE FACTOR 2* (*ERF2*), were upregulated in *tdr4-cr1* and *mbp7-cr1* (Fig. 2D). In addition, the expression levels of genes contributing to ABA signal transduction, including genes encoding the ABA receptor pyrabactin resistance 1-like (PYL; e.g., PYL4/6/7/8/9/10), were upregulated in *tdr4-cr1* and *mbp7-cr1* (Fig. 2E). These findings suggest that the effects of TDR4 and MBP7 on ethylene biosynthesis and signal transduction differ from those of RIN (Bemer et al., 2012). This possibility was supported by the expression patterns of cell wall-related genes (Fig. 2G). Distinct auxin response gene expression patterns were detected in *tdr4-cr1* and *mbp7-cr1* (Fig. 2F). Moreover, *mbp7-cr1* produced smaller fruit than *tdr4-cr1* and AC (Fig. 1A). We speculated that the smaller fruit of *mbp7-cr1* may be related to differences in ARFs, but this possibility will need to be experimentally verified (Fenn and Giovannoni, 2021). Similar to the results of previous studies, we revealed the partially redundant effects of TDR4 and MBP7 on fruit ripening, with MBP7 also modulating early fruit growth and development (Wang et al., 2014; Wang et al., 2019).

### The expression of the regulatory genes controlling carotenoid and ABA accumulation is mediated by ripening-related TFs in tomato

Carotenoids are important dietary nutrients for humans. For example, lutein and zeaxanthin have beneficial effects on eyesight, whereas α-carotene and β-carotene possess antioxidant properties. To clarify the mechanism underlying the color differences between the mutant and AC tomato fruit, we analyzed the expression levels of genes involved in carotenoid biosynthesis in tomato (Fig. 3) (Liu et al., 2015).

**Fig. 3.**
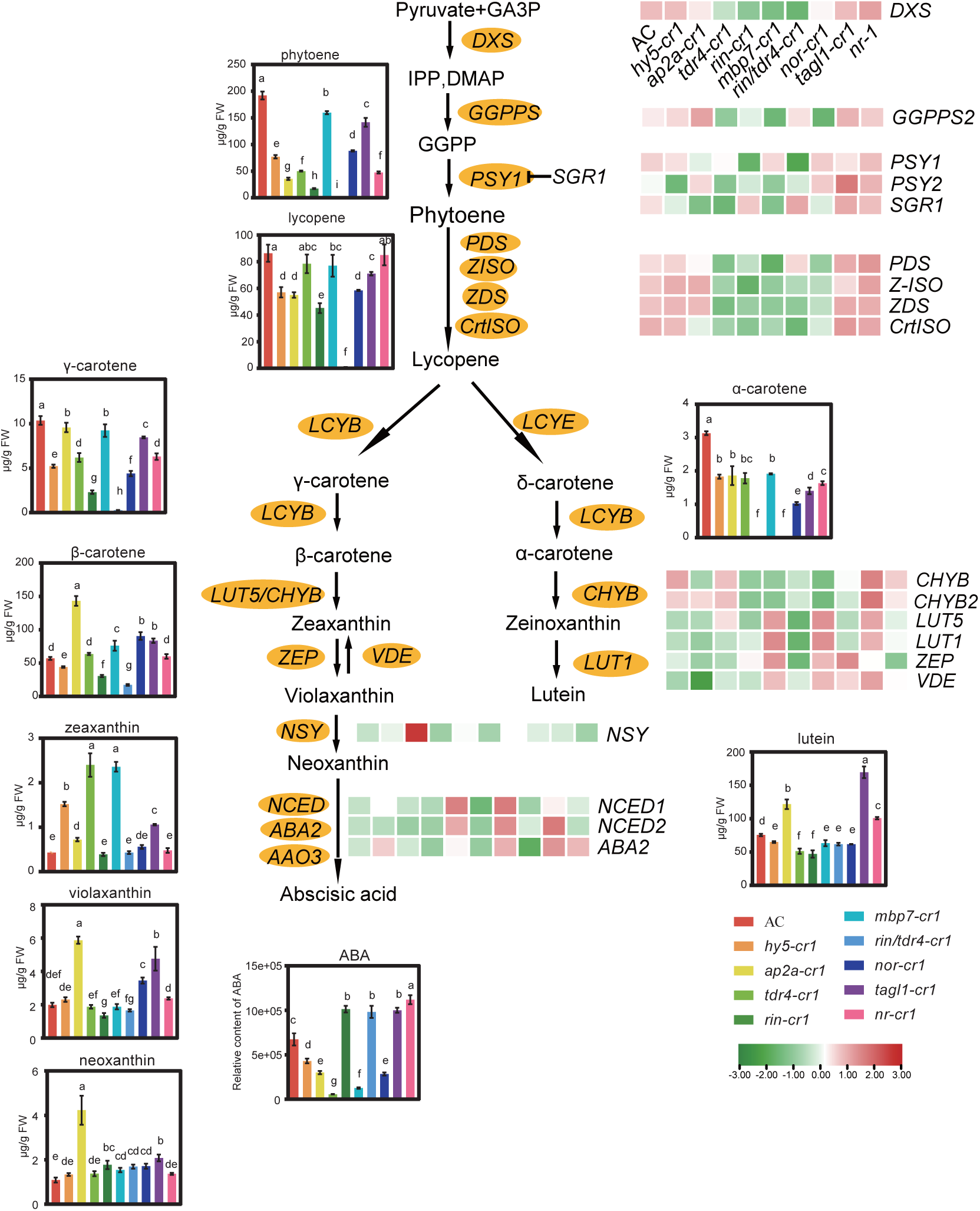
Schematic representation of carotenoid biosynthesis in tomato. The carotenoid contents and relative ABA contents in the ripe mutant and AC tomato fruit are presented in boxes. Data are presented as the mean ± SD (n = 3). Different letters indicate significant differences (P < 0.05) relative to the AC values. Expression patterns of genes involved in carotenoid metabolism in nine mutants and AC. Red color indicates highest expression, and green indicates lowest expression.

Interestingly, the expression patterns of the carotenoid biosynthesis-related genes in *hy5-cr1*, *ap2a-cr1*, and *nr-cr1* were similar to those in the WT. In the *hy5-cr1* mutant, there was a significant decrease in the carotenoid content following the 3-fold decrease in the transcription of the β-carotene hydroxylase gene *CHYB*, but there was only a slight decrease in the expression of *PSY1*, which is the primary gene responsible for carotenoid biosynthesis (Fray and Grierson, 1993). In the *ap2a-cr1* mutant, the total carotenoid content decreased significantly, but there were notable increases in the β-carotene, violaxanthin, neoxanthin and lutein contents. The transcription of both *Geranylgeranyl diphosphate synthase* (*GGPPS2*) and *beta-ring hydroxylase* (*LUT5*) increased significantly, which was in contrast to the slight decrease in the *PSY1* expression level. Moreover, the expression of *STAY-GREEN* (*SGR1*), which promotes chlorophyll degradation, was downregulated by approximately 10-fold in *ap2a-cr1*. In *tagl1-cr1*, the *PSY2*, *CHYB*, *LUT5*, and epsilon-ring hydroxylase (*LUT1*) expression levels were upregulated (relative to the corresponding levels in AC), indicating that TAGL1 may promote lutein accumulation and lycopene degradation via the upregulated expression of *CHYB*, *LUT5*, and *LUT1*. These findings suggest that *AP2a* and *TAGL1* may be potential targets for optimizing the contents of specific carotenoids in tomato fruit. The *RIN*, *TDR4*, and *MBP7* genes encode MADS-box TFs that function as important regulators of carotenoid synthesis. The *DXS*, *GGPPS2*, *PSY2*, *phytoene desaturase* (*PDS*), *ζ-carotene isomerase* (*Z-ISO*), *ZDS*, *carotenoid isomerase* (*CrtISO*), and *CHYB* transcript levels decreased significantly in *rin-cr1*, *rin/tdr4-cr1*, *tdr4-cr1*, and *mbp7-cr1*, indicative of an overlap between the targets of RIN, MBP7, and TDR4 (Bemer et al., 2012). The expression of *PSY1*, which is involved in the production of phytoene and lycopene, decreased by approximately 5.62-fold in *rin-cr1* and by 6.49-fold in *rin/tdr4-cr1* (compared with the WT level); this gene is reportedly targeted by RIN and TDR4 (Martel et al., 2011; Fujisawa et al., 2014). However, *PSY1* was similarly expressed at a slightly lower level in *tdr4-cr1* and *mbp7-cr1* than in the WT (Bemer et al., 2012). These results confirm that RIN, TDR, and MBP7 can influence carotenoid synthesis by regulating *PSY1* and *SGR1* expression (Fujisawa et al., 2013; Fujisawa et al., 2014). Earlier research showed that NOR can promote the accumulation of carotenoids by binding directly to the *GGPPS2* promoter and activating expression (Jiang et al., 2020). The *GGPPS2* expression level decreased substantially in *nor-cr1*, in which the *PDS*, *Z-ISO*, *ZDS*, and *CrtISO* expression levels also decreased slightly, implying NOR may regulate the expression of these genes associated with carotenoid biosynthesis. Therefore, RIN is a key TF contributing to carotenoid accumulation. The other TFs also affect carotenoid accumulation through their regulatory effects on the expression of genes involved in carotenoid biosynthesis (Giovannoni et al., 2017). The downregulated expression of *SGR1* can increase PSY1 levels as well as the lycopene and carotene contents in red ripe fruit, thereby linking carotenoid metabolism and chlorophyll metabolism. The expression of *SGR1* is positively regulated by RIN, NOR-like1, and TDR4/MBP7 (Fujisawa et al., 2014; Gao et al., 2018). In the present study, the *SGR1* expression level decreased by 15.12-fold in *tdr4-cr1*, 9.25-fold in *mbp7-cr1*, 2.16-fold in *nor-cr1*, and 10.62-fold in *ap2a-cr1*. These results suggest that AP2 and NOR may also regulate *SGR1* expression.

Carotenoids are precursors for ABA synthesis, which is required for fruit ripening (Kai et al., 2019). In the present study, we analyzed the expression of ABA biosynthesis-related genes. The *9-cis-epoxycarotenoid dioxygenase* (*NCED1*) transcript levels were almost 4-times higher in *rin-cr1* and *rin/tdr4-cr1* than in the WT, resulting in increased ABA contents. Silencing *NCED* decreases the endogenous ABA level and increases the ethylene content (Ji et al., 2014). This also suggests that RIN negatively controls ABA production by regulating *NCED* expression. The *short-chain dehydrogenases* (*ABA2*) expression level was upregulated by 1.44-fold in *nr-cr1* and by 1.35-fold in *tagl1-cr1*, while it decreased by 7-fold in *ap2a-cr1*, 3.5-fold in *tdr4-cr1*, and 2.37-fold in *mbp7-cr1*; these changes were consistent with the observed changes in the ABA contents. Accordingly, AP2a, TDR4, and MBP7 may help regulate ABA biosynthesis by controlling the expression of *ABA2* during the fruit ripening process. Our findings indicate that the TFs encoded by these genes modulate the ripening of tomato fruit by regulating the expression of distinct carotenoid-related genes. Accordingly, we speculated that these TFs have diverse effects on the accumulation of certain metabolites.

### Metabolome analyses clarified the effects of these TFs on fruit ripening

To more precisely elucidate how these TFs affect fruit metabolite accumulation, we compared the metabolites between the WT and mutant fruit samples. The method for the targeted metabolomics analysis has been used to identify numerous metabolites in tomato (Li et al., 2020; Yang et al., 2022). The 1,052 metabolites identified in the current study were divided into 12 clusters (Fig. 4A and Supplementary Table S14). Additionally, 900 of these metabolites were detected in all samples (Supplementary Figure S6), suggesting the main differences among the mutants were in terms of metabolite contents rather than metabolite compositions.

**Fig. 4.**
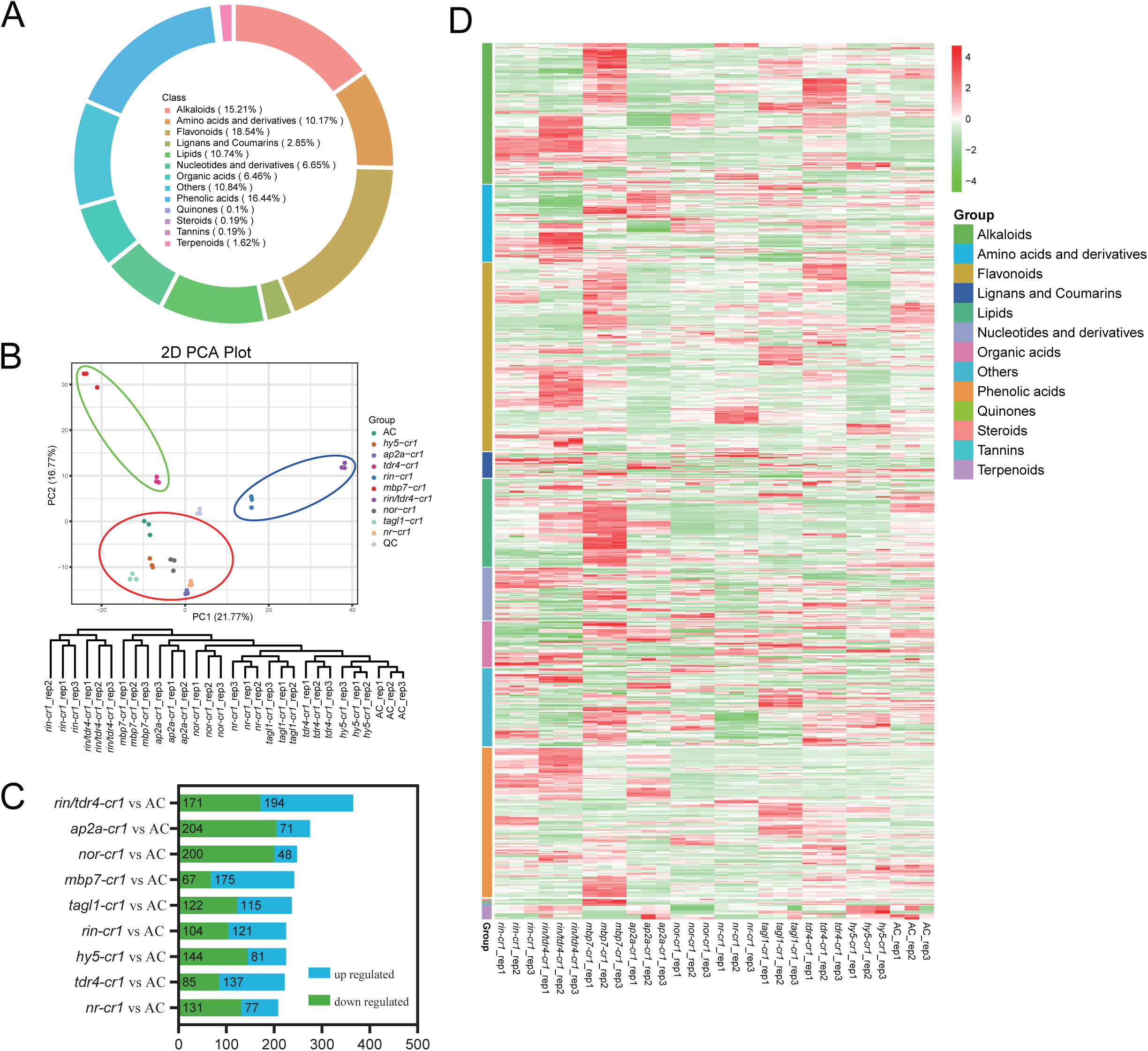
Metabolome analysis of the mutants and AC during the fruit ripening stage. (A Pie chart with metabolite categories. (B) PCA plot and cluster dendrogram of the metabolome data for the fruit ripening stage of nine mutants and AC. (C) Differences in the metabolites identified in the mutants and AC. (D) Overview of the differentially accumulated metabolites between the mutants and AC.

The hierarchical cluster analysis and PCA revealed the clustering of the *hy5-cr1*, *ap2a-cr1*, *tagl1-cr1*, *nr-cr1*, and *nor-cr1* mutants and the WT, indicative of the relatively minor changes to metabolite accumulation induced by the corresponding genetic mutations. In contrast, *rin-cr1* and *rin/tdr4-cr1* were clustered together and *tdr4-cr1* was close to *mbp7-cr1*, indicating the metabolite compositions and contents differed significantly between these four mutants and the WT (Fig. 4B). These findings were consistent with the RNA-seq data (Fig. 2A). A total of 888 differentially accumulated metabolites (DAMs) were detected between the WT and the nine gene-edited tomato mutants (Fig. 4C, Supplementary Figure S7 and Table S15). Among the DAMs, phenolic acids and flavonoids were the major secondary metabolites in fruit (38.9%). The heatmaps of the DAM profiles reflected the diversity between the mutants and AC. Specifically, alkaloid metabolism increased in *mbp7-cr1*, *tdr4-cr1*, and *rin/tdr4-cr1*. In addition, lipid metabolism increased in *mbp7-cr1*, but flavonoid metabolism decreased in *ap2a-cr1* and *hy5-cr1* (Fig. 4D). The enriched KEGG pathways among the DAMs detected by the *hy5-cr1 vs* AC, *ap2a-cr1 vs* AC, and *tagl1-cr1 vs* AC comparisons were phenylpropanoid biosynthesis, flavonoid biosynthesis, and flavone and flavanol biosynthesis. In contrast, linoleic acid metabolism and alpha-linolenic acid were the enriched pathways among the DAMs detected by the *hy5-cr1 vs* AC, *nor-cr1 vs* AC, *nr-cr1 vs* AC, *mbp7-cr1 vs* AC, and *tdr4-cr1 vs* AC comparisons (Supplementary Figure S8). The DAMs revealed by the *rin-cr1 vs* AC and *rin/tdr4-cr1 vs* AC comparisons were mainly associated with propanoate metabolism, caffeine metabolism, phenylpropanoid biosynthesis, and flavone and flavanol biosynthesis. The biosynthesis of various alkaloids was an enriched pathway among the DAMs detected by the *tdr4-cr1 vs* AC and *rin/tdr4-cr1 vs* AC comparisons (Supplementary Figure S8).

### Transcriptional regulation of the metabolite biosynthetic genes in tomato

Tomato fruit are rich in phenolic compounds, carotenoids, and vitamin C, which have nutritional and medicinal effects on humans (Chaudhary et al., 2018). Increases in the carotenoid and flavonoid contents are accompanied by a significant decrease in the α- tomatine content during fruit ripening (Muir et al., 2001; Iijima et al., 2009). To assess the effect of TFs on phenolic components, we screened for phenolic acids and flavonoids synthesized via the phenylpropanoid pathway (Fig. 5A). The predominant phenolic acids in tomato fruit are caffeic acid and chlorogenic acid, whereas the major flavonoids are naringenin chalcone (chalcone) and rutin (flavanol) (Chaudhary et al., 2018). In the present study, knocking out *HY5* resulted in a decrease in the flavonoid contents of fruit (86 of 97 DAMs were downregulated). Among the 86 downregulated metabolites, 21 were associated with flavonoid biosynthesis or flavone and flavanol biosynthesis (including rutin) (Fig. 5B and Supplementary Table S16). Moreover, the *chalcone synthase* (*CHS1*), *CHS2*, flavanone 3-hydroxylase (*F3H*), and *flavonol synthase* (*FLS1*) expression levels decreased significantly (Fig. 5C), indicating that HY5 may regulate the expression of biosynthetic genes to control flavonoid accumulation (Zhang et al., 2022). Of the 85 differentially accumulated flavonoids, 81 were significantly less abundant in *ap2a-cr1* than in the WT, including 12 metabolites related to flavonoid biosynthesis. In contrast, the contents of six phenolic acids involved in phenylpropanoid biosynthesis (including caffeic acid, *p*-coumaroyl shikimic acid, *p*-coumaroyl quinic acid, *p*-coumaraldehyde, coniferin, and syringin) increased significantly in *ap2a-cr1* (Fig. 5D and Supplementary Table S17). The RNA-seq analysis detected genes involved in phenolic acid biosynthesis with markedly increased expression levels, including *phenylalanine ammonia-lyase* (*PAL*), *cinnamoyl-CoA reductase* (*CCR*), *cinnamyl-alcohol dehydrogenase* (*CAD*), *caffeoyl shikimic acid esterase* (*CSE*), and *shikimate O-hydroxycinnamoyltransferase* (*HCT*). In particular, the *PAL6* and *CCR* expression levels were respectively 26.51-fold and 7.01-fold higher in *ap2a-cr1* than in the WT. In contrast, the expression levels of the two *beta-glucosidase* (*BGL)* genes *BGL1* (Solyc01g010160) and *BGL2* (Solyc01g010170) decreased by 63-fold and 36-fold, respectively, suggesting the transcription of these genes is regulated by AP2a (Fig. 5E). Knocking out *TDR4* and *MBP7* caused the fruit flavonoid levels to increase (relative to the WT levels). These observations were in accordance with the findings of a previous study that showed *TDR4* and *MBP7* affect flavonoid accumulation in tomato (Fujisawa et al., 2014). In the *tagl1-cr1* fruit, there was a considerable decrease in the contents of flavonoids, including rutin, which is similar to the results of an earlier study that indicated the overexpression of *TAGL1* promotes flavonoid accumulation (Itkin et al., 2009).

**Fig. 5.**
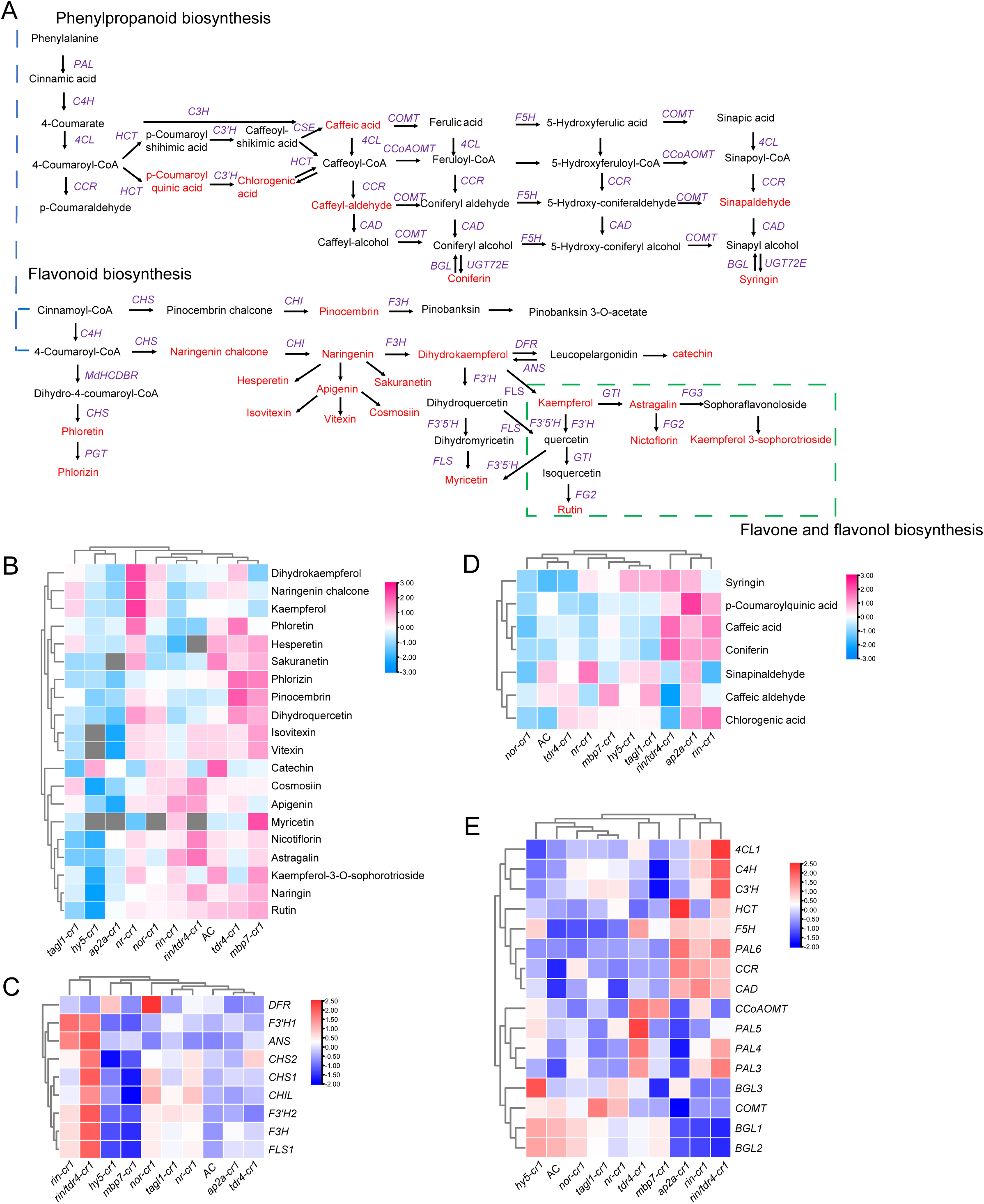
Phenolic acid and flavonoid contents in the mutant and wild-type tomato fruit. (A) Phenolic acid and flavonoid synthesis pathways in tomato. The metabolites that were detected in this study are indicated in red. (B, C) Heatmaps of the DAMs (B) and DEGs (C) related to the flavonoid, flavone, and flavanol synthesis pathways among the mutants and AC. (D, E) Heatmaps of the DAMs (D) and DEGs (E) related to the phenolic acid synthesis pathway among the mutants and AC. The gray block indicates the metabolite was undetectable. The metabolite and transcript data in each row were normalized and standardized.

Linoleic acids (18:2) along with α-linolenic acids (18:3), which are the most abundant polyunsaturated fatty acids in tomato, are essential fatty acids that cannot be synthesized by humans. Moreover, they have a wide range of pharmacological properties potentially useful for treating metabolic syndrome, cancer, inflammation, oxidative stress, and obesity, making them economically important crop components (Hu et al., 2014; Yuan et al., 2022). The KEGG enrichment analysis showed that linoleic acid and α-linolenic acid were enriched among the DAMs detected by the *nr-cr1 vs* AC, *nor-cr1 vs* AC, *tdr4-cr1 vs* AC, and *mbp7-cr1 vs* AC comparisons (Supplementary Figure S8). Linoleic acid and α-linolenic acid metabolites accumulated significantly in *mbp7-cr1*, whereas their contents decreased in *hy5-cr1*, *nor-cr1*, and *nr-cr1* (Fig. 6B). We analyzed the expression levels of the genes associated with linoleic acid and α-linolenic acid metabolism. The *fatty acid desaturase* (*FAD7*) expression level decreased by 4.79-fold in *nor-cr1*, 2.14-fold in *tdr4-cr1*, and 2.03-fold in *hy5-cr1*. These gene expression changes can lead to the desaturation of linoleic acid to produce α-linolenic acid. However, the overexpression of this gene can increase the 18:3/18:2 fatty acid ratio in fruit (Domínguez et al., 2010). In addition, *lipoxygenase* (*LOX*)gene expression also varied in these mutants. The silencing of five *LOX* genes can increase the α-linolenic and linoleic acid contents in tomato fruit (Hu et al., 2014). The *LOXA* and *LOXC* expression levels were upregulated by approximately 4.70-fold and 4.12-fold, respectively, in *nr-cr1*. These results indicate that NOR, TDR4, and HY5 can increase *FAD7* expression, while NR can negatively regulate *LOXA* and *LOXC* expression, which may lead to the accumulation of α-linolenic acids. The transcription of *LOXA*, *LOXB*, and *LOXC* was significantly downregulated in *mbp7-cr1*, suggesting that MBP7 can regulate *LOX* expression, thereby enhancing the accumulation of α-linolenic and linoleic acids. These results further confirmed that two homologous proteins, TDR4 and MBP7, have distinct regulatory effects on linoleic acid metabolism (Fig. 6B).

**Fig. 6.**
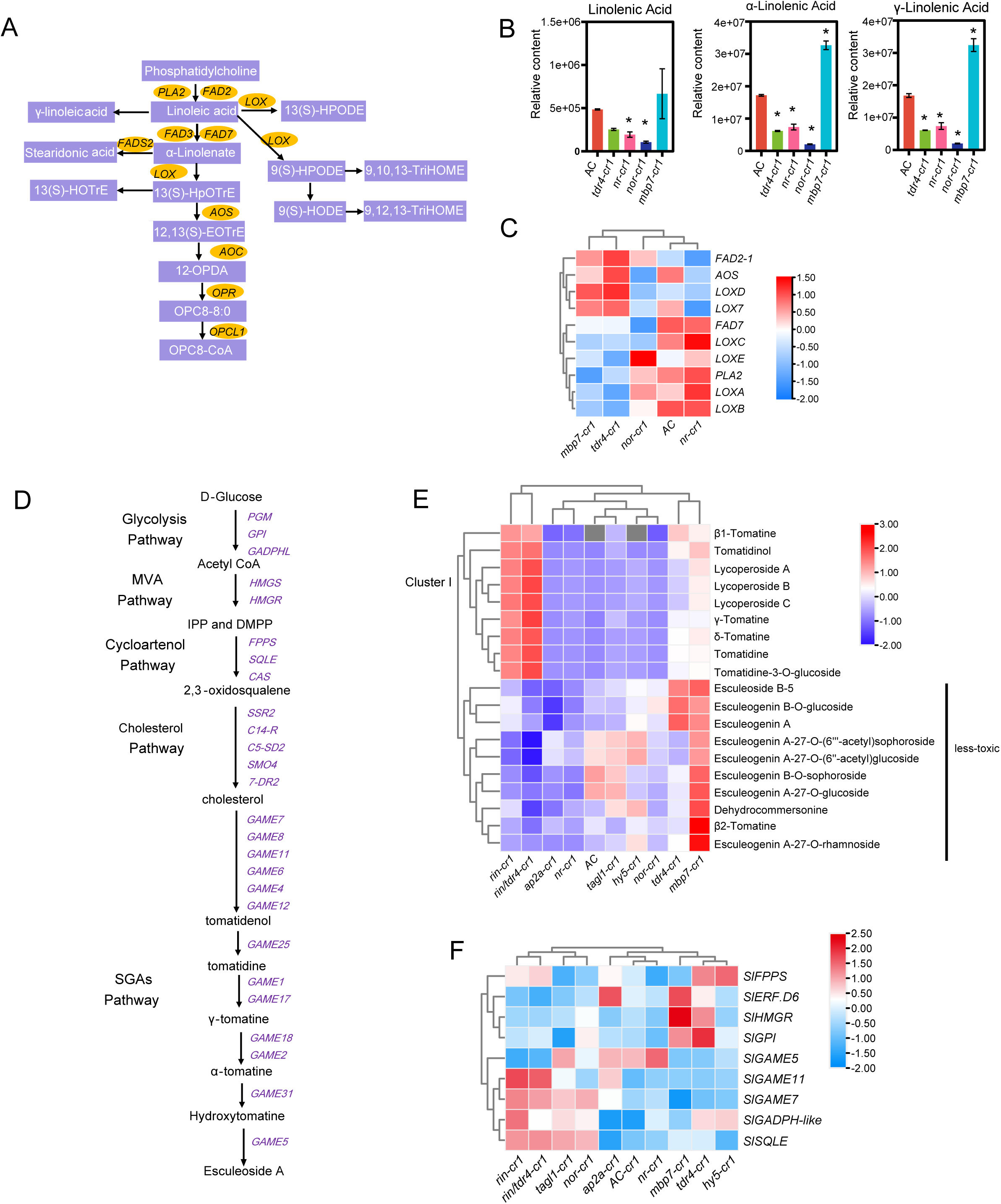
Overview of linoleic acid metabolism and steroidal glycoalkaloid biosynthesis in mutant tomato fruit. (A) Schematic representation of linoleic acid metabolism in tomato. (B) Relative linolenic acid, α-linolenic acid and γ-linolenic acid contents in four mutants and AC tomato fruit. Data are presented as the mean ± SD (n = 3). Asterisks indicate significant differences (*P < 0.05). (C) Expression patterns of linoleic acid metabolism-related genes in four mutants and AC. Gene expression data were normalized to −2 to 1.5. (D) Schematic representation of glycoalkaloid biosynthesis and regulation in tomato. Heatmap of the DAMs (E) and DEGs (F) related to glycoalkaloid biosynthesis in nine mutants and AC. The metabolite and transcript data in each row were normalized and standardized.

Steroidal glycoalkaloids, which are the glycosylated forms of steroidal alkaloids, are metabolites specific to solanaceous plant species (Cárdenas et al., 2016), wherein they protect against diverse natural enemies (Itkin et al., 2013; Paudel et al., 2017; Cárdenas et al., 2019). Some SGAs have a range of pharmacological properties potentially useful for treating various human diseases (Bailly, 2021). A total of 34 SGAs were identified in tomato fruit. The contents of 26, 22, 21, and 15 of these SGAs increased substantially in *rin/tdr4-cr1*, *rin-cr1*, *mbp7-cr1*, and *tdr4-cr1*, respectively. We also observed that most of the 26 DAMs in *rin/tdr4-cr1* and 22 DAMs in *rin* were upregulated. There were no significant differences in the SGA levels between the *tagl1-cr1*, *nr-cr1*, and *hy5-cr1* mutants and AC (Fig. 6D). Both *GLYCOALKALOID METABOLISM* (*GAME*) and the TF gene *ERF.D6* are involved in SGA biosynthesis (Itkin et al., 2013; Cárdenas et al., 2016; Guo et al., 2022). The analysis of these genes and the related pathways indicated that the expression levels of only nine DEGs, including *GAME7*, *GAME11*, and *GAME31*, were significantly upregulated in *rin-cr1* and *rin/tdr4-cr1*, while *ERF.D6* expression was markedly upregulated in *tdr4-cr1* and *mbp7-cr1* (relative to the corresponding AC levels) (Fig. 6E). These results imply that RIN, MBP7, and TDR4 likely regulate the expression of distinct genes in SGA biosynthesis-related pathways (Fig. 6D).

### HY5 helps mediate the accumulation of sugars in tomato fruit

Sugar is an important nutritional component and determines fruit quality. The main soluble sugars in ripening tomato fruit are fructose and glucose. Our metabolomics data showed that the glucose content was 30% higher in *hy5-cr1* than in the WT (Fig. 7A). We also analyzed the sugar content using a high-performance liquid chromatography and mass spectrometry (HPLC-MS) system. The fructose and glucose contents were significantly higher in two independent *HY5* knockout lines than in the WT (Fig. 7B–D). To further clarify how HY5 regulates sugar accumulation, we examined the genes potentially involved in sucrose metabolism as well as the downregulated genes detected by the *hy5-cr1 vs* AC comparison (Supplementary Figure S9 and Table S8). The RNA-seq analysis indicated that the *SuSy* (Solyc12g009300) expression level was 2.29-fold higher in *hy5-cr1* than in the WT. Notably, the expression of *SWEET12c* (Solyc05g024260), which encodes a sugar will eventually be exported transporter (SWEET) (Carrari et al.), was downregulated by 9.27-fold in the *hy5-cr1* fruit (relative to the expression level in the AC fruit) (Fig. 7E). The SWEET proteins reportedly involved in the transport of sugars in tomato have been divided into four distinct clades (Eom et al., 2015; Feng et al., 2015; Ko et al., 2021; Zhang et al., 2021). For example, SWEET12c, which belongs to clade III, is a sucrose and hexose transporter (Eom et al., 2015). The genes encoding other SWEETs were expressed at very low levels during the red ripening stage in the *hy5-cr1* and WT fruit (Supplementary Figure S10). A previous study confirmed that overexpressing *SWEET1a* can decrease the glucose level, resulting in an altered hexose composition in ripening tomato fruit (Shammai et al., 2018). Conversely, the silencing of *SWEET7a* and *SWEET14* via RNAi can increase the fructose and glucose contents in ripening tomato fruit (Chen et al., 2010; Zhang et al., 2021). In the present study, the *SWEET12c* transcript level and hexose content were respectively lower and higher in the *hy5-cr1* tomato fruit than in the WT fruit. To determine whether HY5 directly regulates *SWEET12c* transcription during the ripening of tomato fruit, we analyzed the *SWEET12c* promoter, which revealed three G-box-binding sites (G-box 1: GACGTA-motif; G-box 2: TACGTA-motif; and G-box 3: CACGTT-motif). The ability of HY5 to bind directly to the *SWEET12c* promoter was assessed by conducting a yeast one-hybrid (Y1H) assay and an electrophoretic mobility shift assay (EMSA). The assay results showed that HY5 can bind directly to the G-box element (CACGTT-motif) of the *SWEET12c* promoter (Fig. 7F). Moreover, dual-luciferase assays demonstrated that HY5 can activate *SWEET12c* transcription (Fig. 7G and H). In *Arabidopsis thaliana*, HY5 modulates sucrose metabolism by binding to the *SWEET11* and *SWEET12* promoters to activate transcription (Chen et al., 2012; Chen et al., 2016). Hence, in tomato, HY5 increases *SWEET12c* expression and regulates sucrose accumulation (Fig. 7I). Our findings suggest that a loss-of-function mutation to HY5 and the subsequent decrease in *SWEET12c* expression may increase the tomato fruit sugar content. Specifically, HY5 controls the expression of RRGs involved in various metabolic pathways, such as the sugar transport and storage, phenylpropanoid, and carotenoid pathways, independent of ethylene and ABA. The findings of this study imply that suppressing the expression of *HY5* and *SWEET12c* in tomato plants may increase the fruit sugar content without altering the conversion of SGAs (e.g., α-tomatine) to less toxic metabolites (Fig. 6E and Fig. 7C). The study data may be useful for improving the quality of tomato products, with positive implications for the tomato industry.

**Fig. 7.**
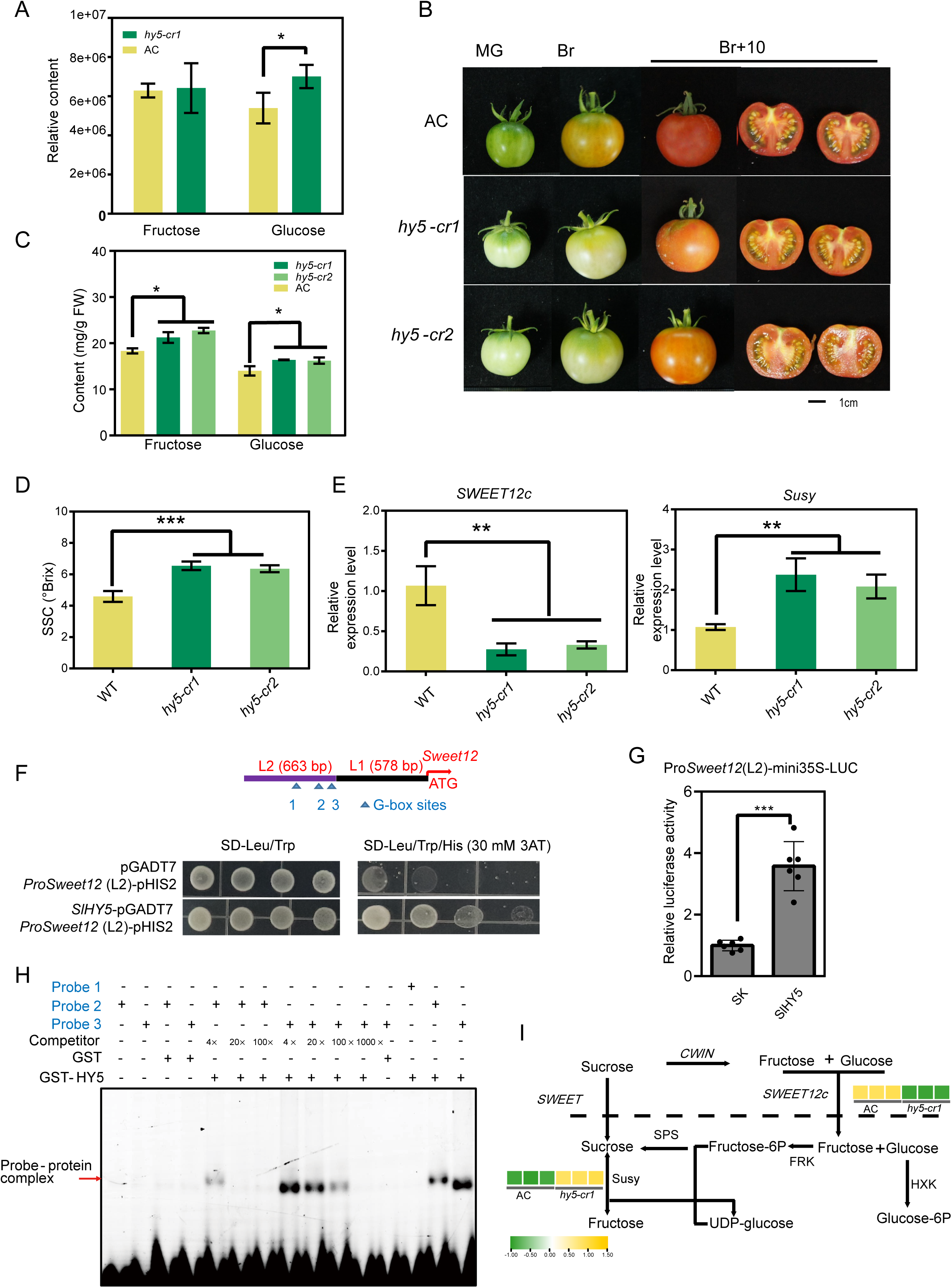
HY5 activates *SWEET12c* expression to regulate sugar accumulation in tomato fruit. (A) Relative fructose and glucose contents in *hy5-cr1* and AC tomato fruit. Data are presented as the mean ± SD (n = 3). Asterisks indicate significant differences (*P < 0.05). (B) Phenotypes of the *hy5* mutant and AC tomato fruit at different development stages. The *hy5-cr1*, *hy5-cr2*, and wild-type fruit at the mature green (25 DPA), break (38 DPA), and ripening (46 DPA) stages are presented. (C) Fructose and glucose contents of the *hy5* mutants and AC at 46 DPA. Data are presented as the mean ± SD (n = 3). Asterisks indicate significant differences (*P < 0.05). (D) Soluble solids content (SSC) of the *hy5* mutants and AC at 46 DPA. Data are presented as the mean ± SD (n = 3). Asterisks indicate significant differences (***P < 0.001). (E) Relative *Susy* and *SWEET12c* transcript levels in fruit at 46 DPA. Values are expressed relative to the wild-type level. Data are presented as the mean ± SD (n = 3). Asterisks indicate significant differences (**P < 0.01). (F) A Y1H assay indicated that HY5 can bind directly to the *SWEET12c* promoter containing the G-box motif. The arrow represents the C/G-box motif. (G) A dual-luciferase reporter assay verified that HY5 affects the *SWEET12c* promoter activity. Error bars represent the SD (n = 6). Asterisks indicate significant differences (two-tailed Student’s *t*-test; ***P < 0.001). (H) The EMSA results reflected the ability of HY5- GST to bind directly to the *SWEET12c* promoter. Lanes 3 and 4 represent the negative control (GST incubated with the labeled probe). Unlabeled probes (4-, 20-, 100-, and 1,000-fold excess) were used as competitors. The probes used for the EMSA were Probe 1 (G-box 1), Probe 2 (G-box 2), and Probe 3 (G-box 3), which were designed and labeled with Cy5. (I) Expression patterns of *SWEET12c* and *Susy*, which influence sucrose metabolism in tomato.

## Conclusion

In this study, the CRISPR/Cas9 system was used to generate mutants. The targeting of fruit ripening-related TF genes resulted in fruit with various colors and distinct carotenoid contents. Although some mutants (e.g., *mbp7-cr1* and *tagl1-cr1*) produced fruit with a similar appearance, there were differences in the carotenoid contents. The comparison of metabolite and transcript profiles between the mutant and WT fruit indicated that the analyzed TFs have distinct regulatory roles affecting metabolite accumulation. Specifically, HY5, AP2a, and TAGL1 can increase flavonoid contents, whereas AP2a inhibits the accumulation of phenolic acids. Linoleic acid and α-linolenic acid metabolism increased substantially in the *mbp7* mutant, but decreased in the *hy5*, *nr*, *nor*, and *tdr4* mutants (compared with the AC levels). The *RIN*, *TDR4*, and *MBP7* genes encode proteins that promote the conversion of SGAs to less toxic metabolites. In the *hy5* fruit, the fructose and glucose contents increased, but pigment synthesis was inhibited, suggesting that HY5 may regulate the balance between sugar and carotenoid contents through a light signaling pathway. The expression of the sucrose metabolism-related gene *SWEET12c* decreased in the *hy5* mutant. According to the Y1H, EMSA, and dual-luciferase assays, HY5 can bind directly to the *SWEET12c* promoter to increase fructose and glucose accumulation. Our study provides genetic information and material useful for creating tomato varieties that produce fruit with ideal colors and high sugar contents.

## Materials and methods

### Plant materials and growth conditions

Tomato (*Solanum lycopersicum*) ‘Ailsa Craig’ (AC) was used as the WT control. The following mutants were generated using the CRISPR/Cas9 system as previously described (Niu et al., 2020): *hy5*-*cr1*, *hy5*-*cr2*, *ap2a*-*cr1*, *rin*-*cr1*, *tagl1*-*cr1*, *tdr4*-*cr1*, *rin*/*tdr4*-*cr1*, *mbp7*-*cr1*, *nor-cr1*, and *nr*-*cr1*. The primers used for constructing vectors and analyzing mutations are listed in Table S18. Plants were grown in a glasshouse with a 16-h day (25 °C)/8-h night (22 °C) cycle and 60% relative humidity. Homozygous *rin*-*cr1*, *tdr4*-*cr1*, *rin*/*tdr4*-*cr1*, and *nor-cr1* lines were produced in a previous study (Niu et al., 2022). Fruits were harvested from each genotype at 46 DPA. Samples from two individual plants were pooled as one biological replicate and immediately frozen in liquid nitrogen before being stored at −80 °C. Three biological replicates were used for the transcriptome and metabolome analyses. The soluble solids content (°Brix) was measured using a handheld digital refractometer (model PR-101; Atago Co., Tokyo, Japan).

### Ethylene production measurement

Two tomato fruit were weighed and enclosed in a 1-L container at room temperature for 1 h. The headspace gas (1 mL) was analyzed using a gas chromatograph (Agilent Technologies 7890A GC System) to measure ethylene production.

### Carotenoid content measurement

To extract carotenoids, tomato fruit samples (50 mg) were ground to a powder, after which 0.5 mL solution comprising n-hexane:acetone:ethanol (1:1:1, v/v/v) was added to the ground material. The mixture was vortexed for 20 min and then centrifuged (12,000 rpm for 5 min at 4 °C). The supernatant was collected and the residue was re-extracted once. The collected supernatants were combined and evaporated. The residue was dissolved in a MeOH:MTBE (1:1, v/v) solution and then filtered through a 0.22 μm membrane before the LC-MS/MS analysis. The extracts were analyzed using a UPLC-APCI-MS/MS system (UPLC, ExionLC™ AD, https://sciex.com.cn/; MS, Applied Biosystems 6500 Triple Quadrupole, https://sciex.com.cn/) as previously described (Geyer et al., 2004).

### Widely targeted metabolome analysis

The metabolomic analysis was conducted by Wuhan Metware Biotechnology Co., Ltd. (Wuhan, China). Three replicates of fruit from the WT and nine mutants were used for profiling the metabolite contents. Briefly, 50 mg ground tomato fruit powder was dissolved with 1.2 mL 70% methanol solution and then vortexed six times for 30 s at 30 min intervals. After centrifuging the solution (12,000 rpm for 3 min), the extract was filtered through a 0.22 μm membrane and analyzed using a UPLC-ESI-MS/MS system (UPLC, Shimadzu Nexera X2, https://www.shimadzu.com.cn/; MS, Applied Biosystems 6500 QTRAP, http://www.appliedbiosystems.com.cn/). The Agilent SB-C18 column (1.8 µm, 2.1 mm × 100 mm) was used for the UPLC. The mobile phase consisted of solvent A (pure water in 0.1% formic acid) and solvent B (acetonitrile in 0.1% formic acid). For the gradient elution, the starting conditions were 95% A and 5% B. After 9 min, a programmed linear gradient changed the mobile phase composition to 5% A and 95% B, which was held for 1 min. Subsequently, the composition was changed to 95% A and 5.0% B in 1.1 min and then held for 2.9 min. The flow rate was set to 0.35 mL/min and the temperature of the column oven was set to 40 °C. The sample injection volume was 2 μL. The eluate was analyzed using an ESI-triple quadrupole-linear ion trap (QTRAP)-MS system. For the ESI source, the temperature was 500 °C; the ion spray voltage ranged from 5,500 V (positive ion mode) to 4,500 V (negative ion mode); the ion source gas I, gas II, and curtain gas were set at 50, 60, and 25 psi, respectively; and the collision-activated dissociation was high. The metabolites were quantified using the multiple reaction monitoring mode. The following criteria were used to identify DAMs: variable importance in projection ≥1 and fold-change ≥2 or ≤0.5.

### Transcriptome sequencing

Fruit pericarps were collected from WT and mutant plants at 46 DPA for the RNA-seq analysis. Total RNA was isolated using the Quick RNA Isolation Kit (Huayueyang Biotechnology Co., Ltd., Beijing, China). The RNA-seq libraries were constructed using the NEBNext^®^ Ultra™ RNA Library Prep Kit according to the manufacturer’s recommendation for Illumina^®^ (NEB, USA). The library quality was assessed using the Agilent 2100 Bioanalyzer system. Finally, the libraries were sequenced on an Illumina platform (PE150). The raw reads were filtered using fastp (v0.19.3) and the clean reads were aligned to the tomato reference genome (SL3.2) using HISAT (v2.1.0) (Kim et al., 2019). StringTie (v1.3.4d) was used for predicting new genes. FeatureCounts (v1.6.2) (Liao et al., 2019) was used to calculate the number of reads mapped to a gene. The fragments per kilobase of exon per million fragments mapped (FPKM) values were used to normalize the expression levels. The DESeq2 (v1.22.1) program and the following criteria were used to identify DEGs: fold-change ≥2 and P < 0.01 (Anders and Huber, 2010). The TBtools program was used for the GO enrichment and KEGG pathway enrichment analyses of the DEGs, with FDR < 0.05 set as the threshold for determining significantly enriched GO terms and KEGG pathways. The hierarchical cluster analysis was conducted using the R package and TBtools (Chen et al., 2020). For the correlation analysis of the transcriptome and metabolome data (log_2_-transformed data), a Pearson correlation coefficient >0.9 and P < 0.05 were set as the thresholds.

### Sugar content determination

The sugar content was measured using acetonitrile. Briefly, 0.5 g ground tomato fruit material was dissolved in 1 mL 70% acetonitrile solution and vortexed for 30 s. Each sample was centrifuged at 11,000 rpm for 5 min, after which the supernatant was collected and evaporated. The pellet was resuspended in 1 mL deionized water, filtered through a 0.22 μm membrane, and analyzed using an HPLC system. Three biological replicates were prepared for each sample.

### Yeast one-hybrid assay

Yeast one-hybrid assays were performed to assess the interaction between HY5 and the *SWEET12c* promoter. The *SWEET12c* promoter sequence was inserted into the pHIS2 vector. The recombinant vector was linearized prior to the transformation of Y187 yeast cells. The *HY5* coding sequence was cloned into the pGADT7 vector to generate the AD-*HY5* construct. The primers used for cloning the *SWEET12c* promoter and *HY5* coding sequence are listed in Supplementary Table S18. The Y187 yeast cells were transformed with the AD-*HY5* and Pro*SWEET12c*-pHIS2 constructs and then screened on SD/−Leu/−Trp medium containing 30 mM 3-AT in plates.

### Electrophoretic mobility shift assay

The *HY5* coding sequence was cloned into the pGEX-4T-1 vector. The recombinant plasmid was then inserted into *Escherichia coli* strain Rosetta (DE3) cells. The primers used for cloning are listed in Supplementary Table S18. The cells were cultured in LB medium at 37 °C until the OD_600_ reached 0.6. The cell culture was cooled to 16 °C before IPTG was added (0.30 mM final concentration) to induce protein expression. The recombinant protein was purified using glutathione beads. The DNA oligos, which were obtained from Shanghai Sangon Biotech, were labeled at the 5′ end with Cy5. The DNA oligos were diluted with ddH_2_O and then mixed with the purified protein for 20 min at 30 °C. After the incubation, the mixture was electrophoresed in a 6% native polyacrylamide gel using a 0.5× Tris-borate-EDTA buffer for 1.5 h at 4 °C and 100 V. Prior to the electrophoresis, the gel was flushed and pre-electrophoresed for 60 min at 4 °C and 100 V. The fluorescence-labeled DNA in the gel was detected using the ChemiDoc Imager (Bio-Rad).

### Dual-luciferase assay

The dual-luciferase assay was performed as previously described (Niu et al., 2016). The *HY5* coding sequence was inserted into the pGreen II 0029 62-SK (SK) vector, whereas the *SWEET12c* promoter was cloned into the pGreen II 0800-LUC (LUC) vector. *Agrobacterium tumefaciens* GV3101 cells transformed with the SK and LUC constructs were infiltrated into *Nicotiana benthamiana* leaves. The LUC and REN luciferase activities were measured using the Modulus luminometer (Promega, Madison, WI, USA).

## Acknowledgments

This work was supported by the National Key Research and Development Program of China (2022YFD2100100, 2021YFA1300401, and NK2022010301), the National Natural Science Foundation of China (32270367), the Office of Education of Anhui Province for Distinguished Young Scholars (2022AH020061), and the Jiangxi “Double Thousand Plan” (jxsq2020101082).

## Accession numbers

The transcriptome data were deposited in the NCBI Sequence Read Archive (BioProject: PRJNA973565).

## Author contributions

Q.N. and Z.L. conceived and designed the study. H.J. performed the bioinformatics analysis. Y-P.X., Y.D., and Y-H.X. contributed to the molecular work and collection of samples. Z.G. revised the manuscript. H.J., Z.L., and Q.N. interpreted the data and wrote the manuscript. All authors read and approved the final manuscript.

